# First record of ‘tail-belting’ in two species of free-ranging rodents (*Apodemus flavicollis and Apodemus agrarius*): Adaptation to prevent frostbite?

**DOI:** 10.1101/2021.04.12.439224

**Authors:** Rafal Stryjek, Michael H. Parsons, Piotr Bebas

**Affiliations:** Institute of Psychology, Polish Academy of Sciences, Jaracza 1, 00-378, Warsaw, Poland; Department of Biological Sciences, Fordham University, 441 East Fordham Road, Bronx, NY, USA; Department of Animal Physiology, Institute of Functional Biology and Ecology, Faculty of Biology, University of Warsaw, 1 Miecznikowa Str., 02-096 Warsaw, Poland

**Keywords:** Apodemus, heat loss, non-commensal rodents, temperature regulation, thermal adaptation

## Abstract

Rodents are among the most successful mammals because they have the ability to adapt to a broad range of environmental conditions. Here, we present the first record of a hitherto unknown thermal adaptation to low temperatures that repeatedly occurred in two species of non-commensal rodents (*Apodemus flavicollis* and *Apodemus agrarius*) between January 16 and February 11, 2021. The classic rodent literature implies that rodents prevent heat loss via a broad range of behavioral adaptations including sheltering, sitting on their tails, curling into a ball, or huddling with conspecifics. Yet, we have repeatedly observed an undescribed behavior which we refer to as “tail-belting”. The behavior was performed during the lowest temperatures, whereby animals - which were attracted out of their over-wintering burrows for a highly-palatable food reward - lift and curl the tail medially, before resting it on the dorsal, medial rump while feeding or resting between feeding bouts. We documented 115 instances of the tail-belting behavior; 38 in *Apodemus agrarius*, and 77 in *Apodemus flavicollis*. In *A. flavicollis*, this behavior was only observed below −6.9C, and occurred more often than in *A. Agrarius*. The latter only demonstrated the behavior below −9.5C. We further detail the environmental conditions under which the behavior is performed, and provide possible functions. We then set several directions for future research in this area.

## Introduction

The yellow-necked mouse, *Apodemus flavicollis* (Melchior 1834), and the striped field mouse, *Apodemus agrarius* (Pallas 1771), are small non-commensal rodents in the family Muridae. They are common across Eurasia, and when conditions are favorable, reproduce rapidly to form numerous populations (Simeonovska-Nikolova, 2007). Despite the differences in their biology and ecology, both *A. flavicollis* and *A. agrarius* have become succesful in the same environments, including urban and peri-urban areas (Simeonovska-Nikolova, 2007; Vukicevic-Radic et al., 2006). However, unlike *Mus* species and commensal rodents, these species are not dependent on human refuse for food, and are therefore, less likely to be observed close to human residences. Though they are of the same genus, a number of differences including morphological, biochemical, and gene location (rearrangement on chromosomes), demonstrate fundamental differences that undermine their presumed systematic relationship (Filippucci et al., 2002; Hille et al., 2002; Rubtsov et al., 2015).

*A. flavicollis* is more closely related to representatives of *Mus* and *Rattus* than to *A. agrarius* (Filippucci et al., 2002; Hille et al., 2002; Rubtsov et al., 2015). Because of these differences, *A. flavicollis* are sometimes considered within the subgenus *Sylvaemus*, while *Sylvaemus* is occasionally used as the genus instead of *Apodemus* (Filippucci et al., 2002; Martin et al., 2000). Similar to other rodents, both species success is partly due to their ability to adapt to highly-variable environmental conditions (Auffray et al., 2009; Bronson and Pryor, 1983; Kay and Hoekstra, 2008). Indeed, *A. flavicollis* and *A. agrarius* are among the best examples of rodents demonstrating tolerance to a broad range of environmental conditions and thus, comprise populations that are widely-distributed from high to low latitudes of Eurasia. These species first ranged in Europe from the southern areas of Scandinavia through western, central, and Mediterranean areas to the northern coast of Africa. Later, two ranges were formed; the western range, covering the south-east Scandinavia through the central and eastern Europe to northern Balkans and central Asia; and the far eastern range, from the south of Russia through eastern China, including the Pacific coast. As a result, representatives of both species persist in disparate areas across temperate, subtropical, and tropical climates. *A. agrarius* is also found in the continental climate zones where seasonal and daily temperature fluctuations can range from *ca.* 30°C to −30°C.

Given the widespread distribution, these two species could be excellent rodent subjects for studies on adaptations to extreme temperatures. Such previous studies have included efficient mechanisms of thermoregulation studied at the molecular and sub-cellular levels (Klaus et al., 1988), basal metabolic rate and thermogenesis (Bligh et al., 1990; Boratyński and Szafrańska, 2018; Haim et al., 1995), and behavioral mechanisms such as social thermoregulation (IJzerman et al., 2021; Tertil, 1972). In many studies of thermoregulation in endotherms, particular emphasis is given to characteristics of the protruding, exposed parts of the body (Arad et al., 1989; Hester et al., 2015; Raman et al., 1983; Romanovsky et al., 2002; Tattersall et al., 2009). The presence of exposed organs can be a challenge when the ambient temperature drops below thermoneutrality, thus they must have mechanisms to prevent heat loss. This strategy results in vasoconstriction that reduces blood flow and helps retain heat (O’Leary et al., 1985; Tan and Knight, 2018).

Other mechanisms protecting against heat loss are countercurrent heat exchangers closely-spaced vessels, often organized in retes, supplying warm blood to the protruding parts of the body and draining cool blood. This process allows heat to radiate from arterial to venous blood before it reaches the periphery of the protruding organs, where it could be significantly cooled (Scholander and Krog, 1957). This approach occurs in such species as sloths (Scholander and Krog, 1957), cetaceans (Heyning, 2001) and turtles (Davenport et al., 2015). But among rats (*Rattus spp*.), countercurrent heat exchangers are not likely to be involved in preventing the loss of heat from tails, where vasodilation plays a major role (Dawson and Keber, 1979; Young and Dawson, 1982). Similarly, in *Mus musculus*, tails also appear to contribute little to thermoregulation (Škop et al., 2020).

The threats that could result from the destabilization of the body’s temperature balance, can also be mitigated behaviorally. In rodents, behavioral adaptations include changes in foraging behavior in the Degu, *Octodon degus;* Bozinovic et al., 2000) and deer mice, *Peromyscus maniculatus;* Sears et al., 2009). This seems to imply that, for these species, thermoregulatory abilities may actually be more crucial than mitigating threats from predators (Lagos et al., 1995). Avoiding thermal stress may also involve modifying essential life tasks, such as finding resources at different times between day and night (desert woodrat, *Neotoma lepida;* Murray and Smith, 2012) and different seasons (common vole, *Microtis arvolis;* Hoogenboom et al., 1984). For review of the latter, see (Bennie et al., 2014). Behavioral thermoregulation is also associated with the exploitation of various thermal refuges, such as vegetation plant cover (D’Odorico et al., 2012; Pigeon et al., 2016).

A common behavioral adaptation involves constructing burrows leading to an underground nest, and remaining therein during the coldest periods (Amori et al., 1986; Baláž and Ambros, 2012; Stradiotto et al., 2009). This strategy allows for hoarded food reserves to be utilized in the effective shelter, which *A. flavicollis* especially relies on during the winter. In addition, constructing nests near human settlements or in urbanized areas often increase access to crop resources (Balčiauskas et al., 2020) and therefore, increases the chance of survival for *A. agrarius* (Andrzejewski et al., 1978; Gliwicz and Taylor, 2002). For comparisons of both species’ biology see (Baláž and Ambros, 2012).

Another behavioral phenomenon observed at low temperature is curling into a ball-like posture in order to keep warm, and adopting this posture to reduce the surface-to-volume ratio (Terrien et al., 2011). Curling, commonly observed in mammals, including domestic pets, has also been described in the rodent literature (Gosling, 1979; Moinard et al., 1992; Morrison and Tietz, 1957). In many animals, such reductions in body surface area also involve setting the protruding parts of the body so that they adhere to its surface as much as possible (Prestrud, 1991; Scholander, 1955). This behavior is perhaps also important to protect them from damage, such as by frostbite, as the trunk temperature is usually higher and kept relatively constant compared to these protruding parts of the body.

A similar phenomenon, which we now refer to as ‘tail belting’, where the animal lifts and may curl the tail medially, before resting it on the dorsal, medial rump, was observed in both *A. flavicollis* and *A. agrarius,* during feeding and resting between feeding bouts. The behavior occurred when the surrounding temperature was below −6.9°C. in *A. flavicollis,* and −9.5 °C in *A. agrarius*. As far as we know, this phenomenon has not been described in the literature, and here we systematically document the occurrence under particular circumstances.

## Methods and Materials

The observed behavior was recorded during a field study conducted on free-living colonies of yellow-necked mice (*Apodemus flavicollis*) and striped field mice (*Apodemus agrarius*) on a private, suburban property in Warsaw, central Poland (52°20′N 21°03′E, altitude: 80 m). The experiment took place between 1 November, 2020 and 15 March, 2021 during the Winter season. Temperatures during this period ranged from +16°C to −20°C. Based on direct and video observations over five months, we estimate the population size of each colony to be in excess of 10 individuals of each species. Individual recognition was sometimes possible based on distinctive variations in coat patterns, body size, and individual characteristics including marks, scars, wounds, variegation of color and shape of tail.

The study included continuous video recording of two chambers which were constructed to test individual responses to scents from conspecifics and/or predators. Two chambers were constructed from 12mm waterproof plywood and painted with odorless paint. The internal floor dimensions were 35◻×◻40◻cm with a wall height of 70◻cm. Chambers were deployed near cover beside bushes in the natural habitat of neighboring forest and meadows. Malleable bent sewer pipes (70mm diameter and 50cm length; Certus, Cieszyn, Poland) were connected to entrance holes. The bottom of the chambers was covered with 1cm of rinsed sand and replaced twice per week. Animals were baited to the chambers nightly at dusk with 5g of chocolate-nut cream (Nuss Milk Krem; i.e., Nutella; Jackson et al., 2016) placed on 70mm Petri dishes.

For continuous surveillance, we used three infrared cameras (Easycam EC-116-SCH; Naples, FL, USA) connected to a digital video recorder (Easycam EC-7804T; Naples, FL, USA). This setup enabled 24/7 motion detection recording for the duration of the study. Following our repeated observation of this behavior, we also utilized thermal imaging (Seek Thermal Shot SW-AAA thermal camera; Santa Barbara, CA, USA) and recorded two visits to the chambers by yellow-necked mice (both at +1.5°C) and five instances of visits by striped field mice (at −1.5°C, − 2.5°C, 5°C, 6°C, & 8°C).

### Ethics Statement

This observational study was a non-invasive experiment based on the surveillance of free-ranging animals that were free to enter or ignore experimental chambers with food and video cameras. Thus, it did not require permission of the local ethics committee for animal experimentation. The study was carried out on private land with permission of its owners, and all procedures were conducted in accordance with the Polish Animal Protection Act (21 August, 1997).

## Results

We recorded 115 instances of tail-belting (38 in *A. agrarius* and 77 in *A. flavicollis*) during five months of continuous observation of the two colonies. Within this 5-month period, the only instances of tail belting occurred between January 16 and February 11, 2021 during a particularly harsh winter period in Warsaw, Poland (Supplemental Table 1). Given the number of incidents and the colony size of both species, it is possible that over a dozen animals of each species displayed this behavior. While we could not always identify individuals due to their somewhat uniform appearance, we can be certain that at least 8 individuals (4 of each species) displayed this behavior. We were able to distinguish the 8 individuals due to observable differences in coat patterns, body size, and individual characteristics such as scars, or crooked tails.

The lowest chamber temperature recorded during foraging was −17°C for *A. flavicollis* (Figure 1; Supplemental video 1) and −14.5°C for *A. agrarius* (Supplemental video 2). While animals were recorded across many temperatures during the 5-month period, tail-belting was first recorded when the temperature dropped to −6.9°C in *A. flavicollis,* and −9.5°C in *A. agrarius* (Figure 1). Thermal images showed the temperature of the tail equaling dropping well below trunk temperature and, in some cases, equaling ambient temperature below 0°C for both species (Figure 2; Supplemental video 1, 2). The frequency of tail-belting may have increased with additional decreases in temperature (Supplemental Table 1), though we did not quantify this number.

**Figure 1.**
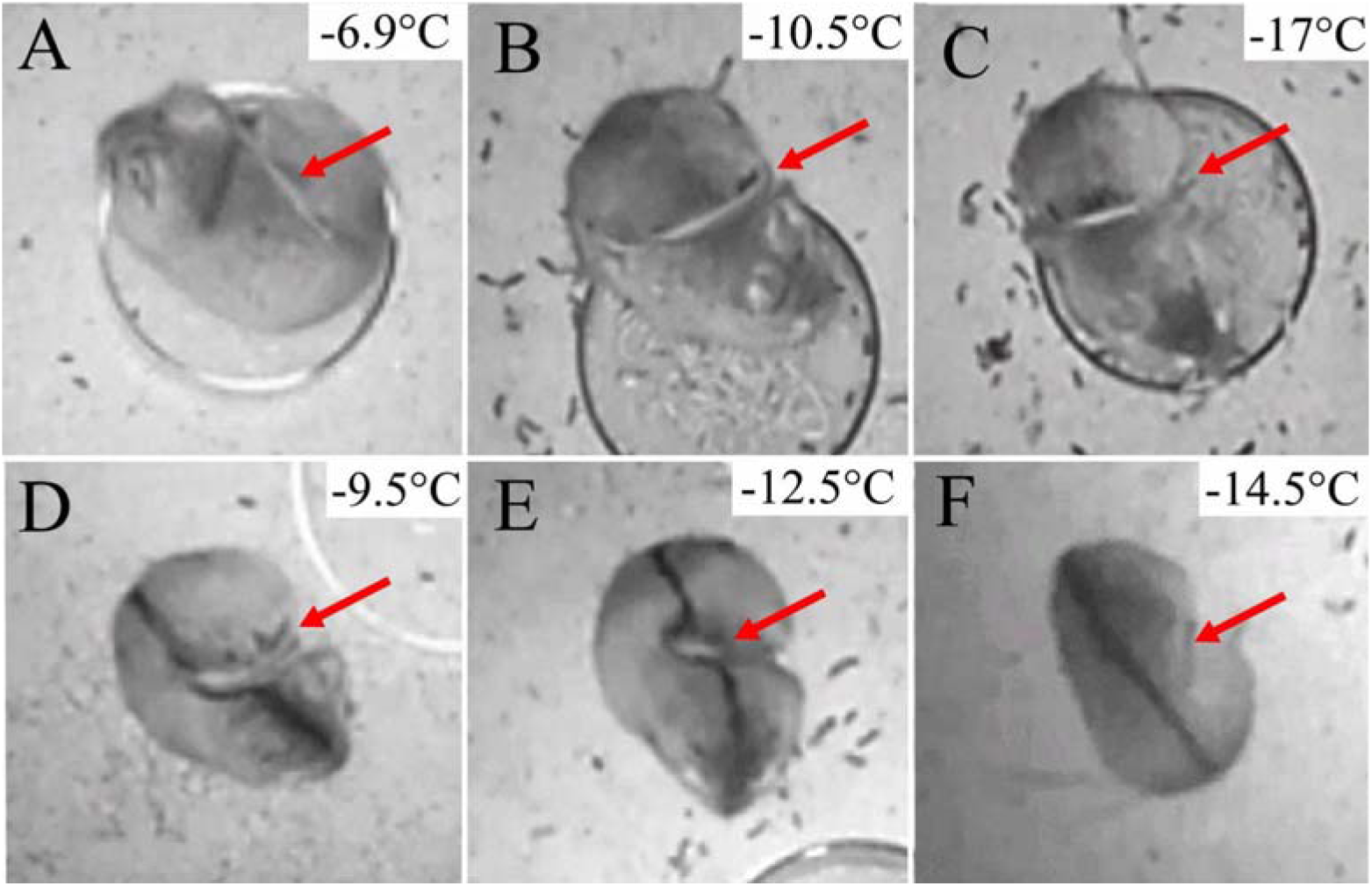
Video stills from IR cameras showing visible tail-belting in mice. A-C: Yellow-necked mice; D-E: Striped field mice. A and D show the lowest temperature that tail-belting was recorded at for each of the two species, C and F show the highest temperature that tail-belting was recorded at for the two species. Red arrows indicate the position of the tail being belted.

**Figure 2.**
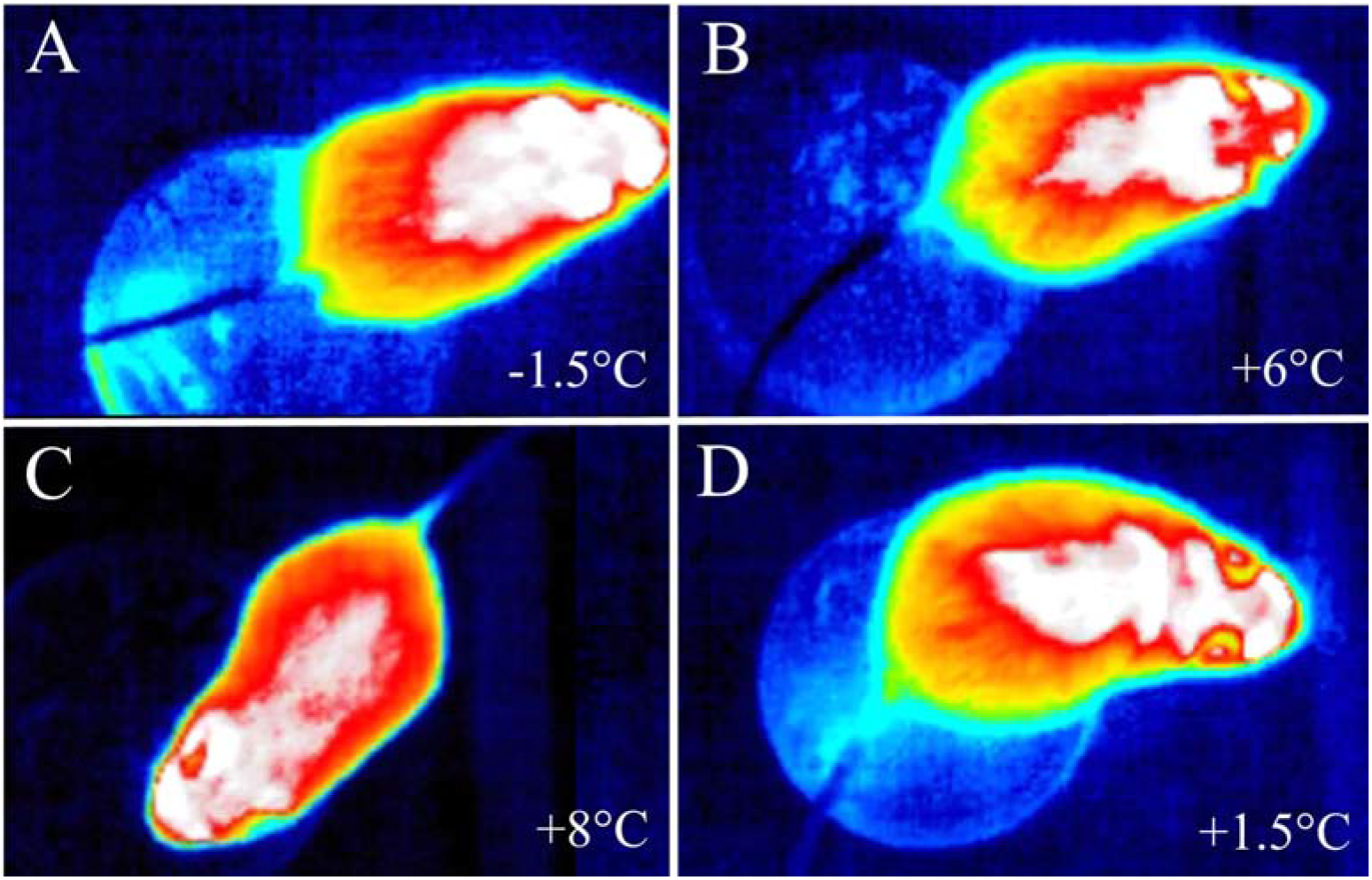
Thermal images showing temperature of the tail equaling ambient temperature (for NETD = 70mK). A-C) a striped field mouse; D) a yellow-necked mouse. NETD= Noise Equivalent Temperature Difference.

## Discussion

We have documented over one hundred instances of tail-belting, a previously undescribed behavior, among two non-commensal rodents. Because this behavior only occurs below a temperature threshold, it is most likely an adaptation to prevent frostbite of an exposed, protruding appendage or freezing of the tail to a surface. Indeed, we assume frostbite prevention, and not decreased heat loss, is the most likely explanation for two reasons. Firstly, thermovision (Figure 2, Supplemental video 1, 2) showed that tails dropped well below trunk temperature and secondly, tails appear to contribute very little to thermoregulation among mice (Škop et al., 2020).

Between the two species, *A. agrarius* demonstrated tail-belting more rarely than *A. flavicollis* and at colder temperatures. This discrepancy is likely because *A. agrarius* has shorter tails and ceased foraging at a higher temperature. Given that rodents are among the most successful and best-known animals, particularly the genus *Mus* (Kay and Hoekstra, 2008), we can only assume this behavior has not previously been documented because of the difficulty of observing small, free-ranging rodent species in sub-optimal conditions in the wild. Additionally, mice minimize foraging in winter while remaining in burrows and consuming hoarded food. Under natural conditions (e.g., without access to our experimental chambers with food), during temperatures when this tail-belting behavior is prominent, most mice would likely not even come out of their burrows. These individuals may have ventured out only because we provided a consistent, aromatic and highly-palatable food on a daily basis for months before and after the cold temperatures of the particularly harsh winter of 2021 in Warsaw, Poland. Indeed, we suspect we only observed this behavior because we set up trials intending to record and assess behaviors in the presence or absence of particular scents near a food reward. Given the relatively distant relatedness between the two species, it is likely that this behavior also occurs in other rodents, particularly free-ranging animals that usually remain inside burrows in colder climates.

Future studies should involve exact measurement of temperature of body parts using thermo-vision cameras, as well as morphological and histological comparisons of the species. Precise linear studies should ensue to determine if decreasing temperatures below the threshold increase the frequency of this behavior. Anecdotally, it appears that this was the case, however, we do not know if temperature is the only factor that causes this behavior. It is unknown whether anyone has reduced the temperature in the laboratory in attempts to induce the behavior in other species. *Mus musculus* is currently the primary model for studying frostbite injuries, however, this is not done by lowering the temperature, but instead by adhering frozen magnets to the skin (Auerbach et al., 2013). Thus, tail-belting would not have been expressed under these conditions. Moving forward, this behavior could also be sought out within laboratory conditions, with direct comparisons between the two *Apodemus* species and *Mus*.

Mice assays are very popular throughout science and account for more than 60% of all laboratory assays with animals used in research in Europe (European Commission, 2020) Thus, there should be many opportunities to examine whether this behavior is also observed among laboratory animals. We do not know if the behavior is an unconditioned reflex, perhaps if this is shown to be the case, then studies could seek to determine a corresponding neural pathway. Regardless of the mechanism, it appears that this behavior is yet another example, among many, of how rodents have become one of the most diverse. adaptable and successful taxa.

## Acknowledgements

This research was self-funded by the authors, with the exception of equipment funds awarded to RS (Polish National Science Centre (NCN) – Grant Number: UMO-2013/09/B/HS6/03435) and PB (Polish National Science Centre (NCN) – Grant Number: UMO-2013/11/B/NZ4/03310). All authors declare no conflict of interest.

## Notes

### Competing Interest Statement

The authors have declared no competing interest.

